# A live-cell marker to visualize the dynamics of stable microtubules

**DOI:** 10.1101/2021.06.23.449589

**Authors:** Klara I. Jansen, Mithila Burute, Lukas C. Kapitein

## Abstract

The microtubule (MT) cytoskeleton underlies processes such as intracellular transport and cell division. Immunolabeling for post-translational modifications of tubulin has revealed the presence of different MT subsets, which are believed to differ in stability and function. Whereas dynamic MTs can readily be studied using live-cell plus-end markers, the dynamics of stable MTs have remained obscure due to a lack of tools to directly visualize these MTs in living cells. Here, we present a live-cell marker to visualize stable MTs and explore their dynamics. We demonstrate that a rigor mutant of kinesin-1 binds selectively to acetylated MTs without affecting MT organization and organelle transport. These MTs are long-lived, do not depolymerize upon nocadozale-treatment or laser-based severing, and display rich dynamics, including undulation, looping and sliding. This marker will help to explore how different MT subsets contribute to cellular organization and transport.

## Introduction

Cells use microtubules (MTs) to establish intracellular organization. These stiff and polarized polymers span long distances throughout cells and serve as tracks for directional transport driven by motor proteins that move towards either plus- or minus-end of MTs. Most MTs switch between phases of polymerization and depolymerization in a process called dynamic instability (Mitchison & Kirschner 1984), which is coupled to the GTP-status of the newly incorporated tubulin subunits at the MT plus-end (Akhmanova & Steinmetz 2015). The exact dynamics of MTs can furthermore be tuned by a wide variety of microtubule-associated proteins (MAPs) that can stabilize or destabilize MTs, either by modulating their end dynamics or by affecting the polymer as a whole. In addition, MTs can undergo post-translational modifications (PTMs) and these are believed to also impact, directly or indirectly, microtubule stability (Janke & Magiera 2020; Roll-Mecak 2020).

Immunocytochemistry on fixed cells stained for different modifications has revealed that the concerted action of these MT-regulating factors can lead to the emergence of distinct, co-existing MT-subpopulations that greatly differ in their chemical composition and stability (Verhey & Gaertig 2007; Burute & Kapitein 2019; Janke & Magiera 2020; Roll-Mecak 2020). Most MTs are composed of tubulin with a C-terminal tyrosine, undergo rapid cycles of growth and shrinkage, and quickly depolymerize upon cold treatment or treatment with MT destabilizing agents. In contrast to these dynamic MTs, stable MTs survive such treatments and are enriched in post-translational modifications, such as C-terminal detyrosination or acetylation. This MT diversity is believed to play an important role in the spatial organization of cells, as different subsets are organized differently and are used by different motor proteins as tracks for organelle transport (Burute & Kapitein 2019). Stable, long-lived MTs polarize towards the leading edge in migrating fibroblasts (Gundersen & Bulinski 1988) and are predominantly localized in the perinuclear area in common cell lines like HeLa, Cos7 and U2OS, whereas dynamic MTs display a more homogenous distribution. In dendrites of hippocampal neurons, stable MTs are enriched in the core of the dendritic shaft, while dynamic MTs mainly localize near the plasma membrane (Tas et al. 2017; Katrukha et al. 2021). In addition, stable and dynamic MTs often display opposite orientations in dendrites (Tas et al. 2017). Furthermore, it has been shown that some motor proteins bind selectively to one subset of MTs. For example, kinesin-1 moves preferentially along stable MTs (Cai et al. 2009; Dunn et al. 2008; Guardia et al. 2016; Tas et al. 2017), whereas Kinesin-3 prefers dynamic MTs (Guardia et al. 2016; Tas et al. 2017; Lipka et al. 2016). The guidance of motors by distinct MT subsets has been implied to underly organelle positioning in non-neuronal cells (Guardia et al. 2016; Mohan et al. 2019; Serra-Marques et al. 2020) and polarized transport in hippocampal neurons (Tas et al. 2017).

Although the functional relevance of MT subsets in the spatial organization of cells is becoming increasingly clear, little is known about the organization and establishment of the MT subsets themselves. How do certain MTs become stabilized and how do they obtain their specific spatial organization? Is the stability of a MT modulated in its ends or along the lattice? And how do PTMs and MAPs relate to MT stability? Studies investigating those questions would greatly benefit from tools that enable direct visualization of MT subsets in living cells. Dynamic MTs can readily be visualized in living cells using plus-end markers or the recently developed tyrosination sensor (Kesarwani et al. 2020), yet this is not the case for stable MTs, which can currently only be visualized using immunocytochemistry. To address this, we set out to develop a live-cell marker for stable MTs to bring this MT subset into focus. After validating that this marker does not perturb the MT cytoskeleton nor organelle transport when expressed at low levels, we explore the dynamics of stable microtubules and the origin of their stability in live cells.

## Results

To visualize stable MTs in living cells, we exploited the intrinsic property of kinesin-1 to move selectively along the subset of stable MTs (Cai et al. 2009; Dunn et al. 2008). Introduction of a G234A rigor mutation perturbs the ATPase activity of this protein (Rice et al. 1999), resulting in a motor protein that has a very low rate of MT unbinding. Similar to the active motor, the G234A rigor mutant of kinesin-1 binds to, and effectively decorates, acetylated MTs (Tas et al. 2017; Guardia et al. 2016; Farías et al. 2017). We reasoned that if we would fuse kinesin-1 rigor to a tandem of the very bright fluorophore mNeongreen (Shaner et al. 2013), we could exploit kinesin-1 rigor as a live-cell marker for stable microtubules (Fig.1A-B and video1). As expected, rigor-2xmNeongreen localized with high selectivity to the subset of acetylated MTs upon expression in U2OS cells (Fig.1C-E). Detyrosination is another PTM that is frequently associated with stable MTs (Janke & Magiera 2020). Accordingly, the subset of detyrosinated MTs is decorated by rigor-2xmNeongreen in U2OS cells (Sup. Fig.1).

**Figure 1.**
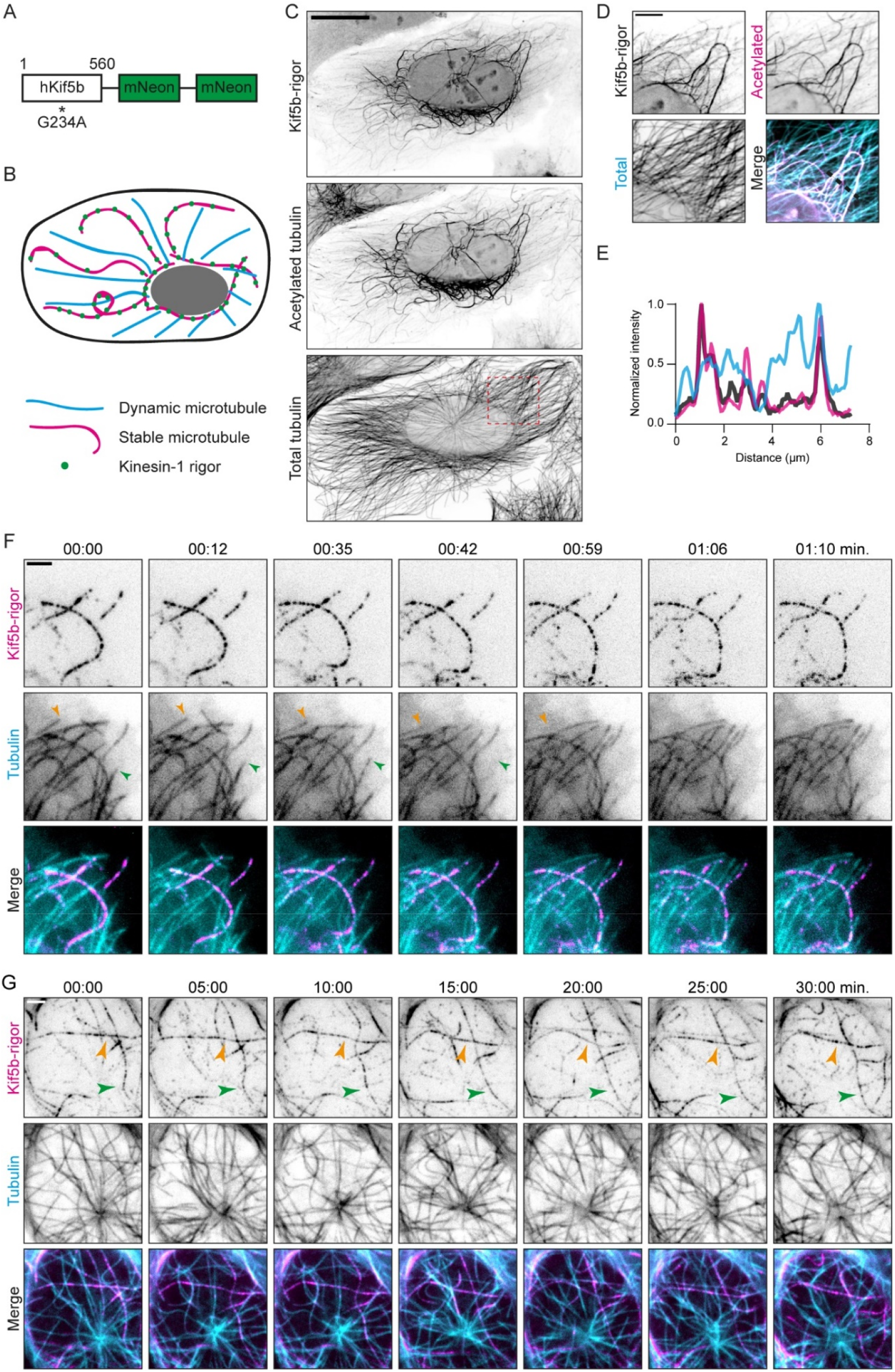
Kinesin-1 rigor as a live-cell marker for stable microtubules. **(A)** Construct: 1-560 truncation of human kinesin-1 containing the G234A rigor mutation fused to a tandem of mNeongreen separated by linker sequences. **(B)** Cartoon: upon expression in cells, rigor-2xmNeongreen decorates stable MTs in a speckle-like fashion. **(C)** Fluorescence images of U2OS cell expressing rigor-2xmNeongreen and immunolabelled for acetylated tubulin and alpha-tubulin shown in inverted contrast. **(D)** Zooms of region indicated by dashed box in (C). **(E)** Intensity profile across the region indicated with dashed line in (D). **(F-G)** Stills from live-cell imaging of rigor-2xmNeongreen and mCherry-tubulin on different time scales (see also Video 2 and 3). Coloured arrows in (F) show examples of MTs displaying dynamic instability. Coloured arrows in (G) indicate rigor-decorated MTs that are retained during the duration of imaging. Scalebars, 20μm (C), 5μm (D), 2.5μm (F-G). Time, minutes:seconds.

To test the potential of rigor-2xmNeongreen as a live-cell marker for stable MTs, we co-transfected U2OS cells with rigor-2xmNeongreen, and mCherry-tubulin to visualize all MTs. Using TIRF-microscopy we observed that many MTs labelled only by mCherry-tubulin displayed dynamic instability, while the MTs decorated by rigor-2xmNeongreen did not, indicating that the MTs marked by rigor-2xmNeongreen are indeed stable on this timescale (1:10 min.) (Fig. 1F). Next, we performed a similar experiment but with an increased imaging duration and we found that some MTs labelled with rigor-2xmNeongreen could be followed for 30 minutes (Fig.1G). In addition, we tested the resistance of rigor-decorated MTs against nocodazole-induced depolymerization. Using Spinning Disk-microscopy, we visualized U2OS cells expressing rigor-2xmNeongreen and mCherry-tubulin. Upon addition of 10μM nocodazole, the majority of MTs labelled by mCherry-tubulin was rapidly lost and the signal became diffusive. In contrast, rigor-MTs remained largely unaffected by the treatment; 30min. after the addition of nocodazole a clear network of rigor-MTs could still be observed (Sup.Fig.2). Together, these data demonstrate that rigor-2xmNeongreen functions as a live-cell marker for stable MTs.

A potential pitfall of our approach to use kinesin-1 rigor as a live-cell marker for stable MTs, is that MT-binding proteins can induce artefacts of the MT cytoskeleton when highly overexpressed. To examine whether kinesin-1 rigor overexpression resulted in microtubule overstabilization, we measured cellular MT acetylation levels as a function of kinesin-1 rigor overexpression levels. We expressed rigor-2xmNeongreen in U2OS cells and performed immunolabeling for acetylated tubulin and alpha-tubulin (Fig.2A). To quantify the effect of rigor-2xmNeongreen expression on MT acetylation levels, we randomly imaged rigor- and non-rigor expressing cells (see Sup. Fig. 1A for example images) and measured and calculated the intensity of acetylated tubulin and rigor-2xmNeongreen normalized to the intensity of alpha-tubulin for individual cells. There was a clear correlation between rigor expression and MT acetylation levels, with high rigor-expressing cells having increased levels of MT acetylation (Fig.2A-B). In addition, high rigor expression induces clear bundling of acetylated MTs. However, at low rigor expression, acetylation levels of rigor-positive cells were within the range of the acetylation levels observed for non-rigor expressing cells (Fig.2A-B; for duplicate see Sup.Fig.1A-B). This indicates that when expressed at sufficiently low levels, rigor-2xmNeongreen does not alter MT acetylation.

**Figure 2.**
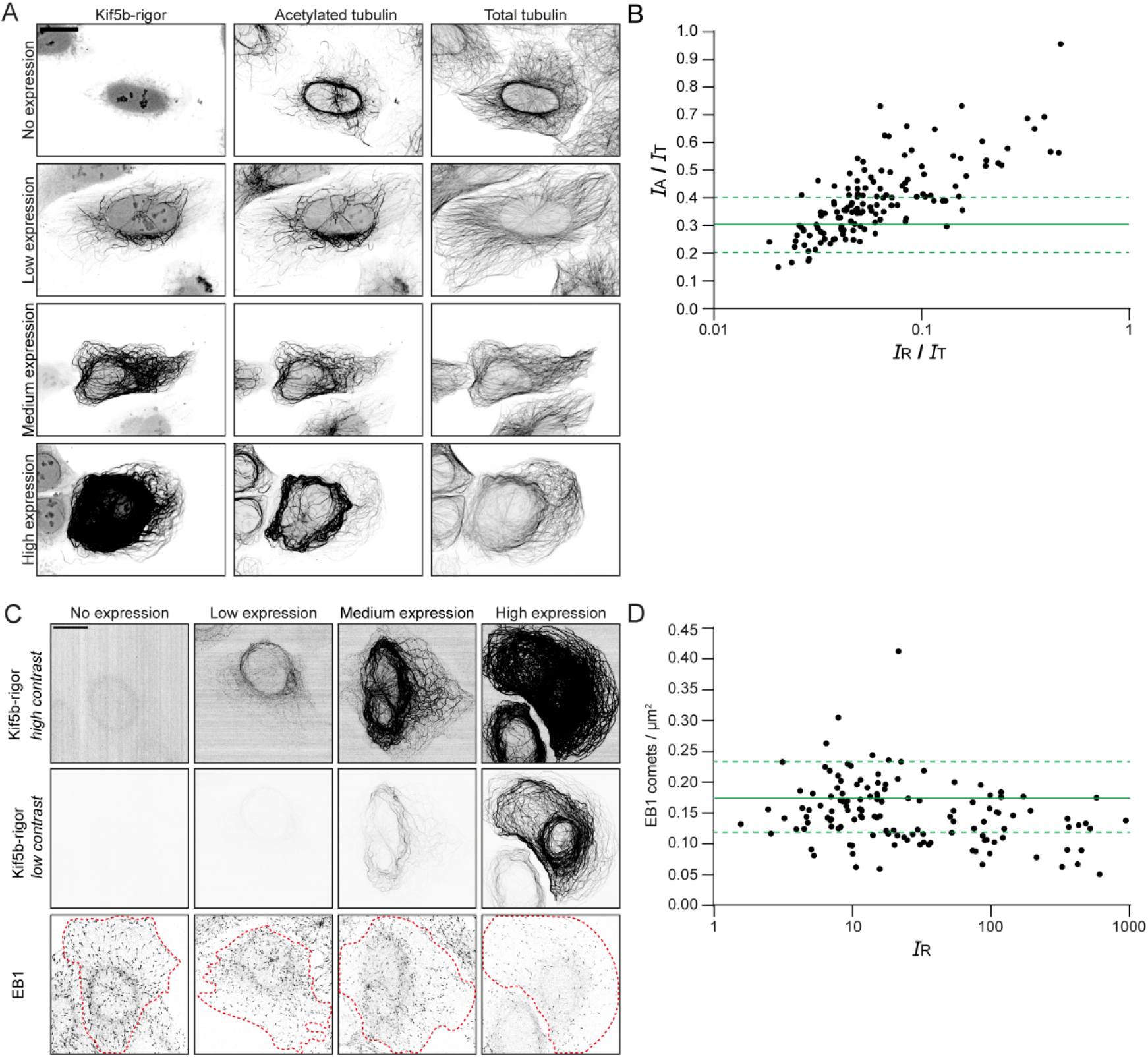
Kinesin-1 rigor has minimal effects on the microtubule cytoskeleton at low expression levels. **(A)** Fluorescence images of U2OS cells stained for acetylated tubulin and alpha-tubulin at different levels of rigor-2xmNeongreen expression. **(B)** Quantification showing the intensity ratio of acetylated tubulin (*I*_A_) over total tubulin (*I*_T_), plotted against the intensity ratio of rigor-2xmNeon (*I*_R_) over total tubulin for individual rigor-expressing cells (each dot represents a single cell, n = 137). Solid green line + dashed lines indicate mean ± SD of the intensity ratio of acetylated tubulin over total tubulin for non-rigor expressing cells (n = 125). **(C)** Fluorescence images of U2OS cells immunolabelled for EB1 at different levels of rigor-2xmNeongreen expression. **(D)** Quantification showing the amount of EB1 comets/μm^2^ for cells with different rigor-2xmNeongreen intensities (each dot represents a single cell, n = 131). Solid green line + dashed lines indicated mean + SD of amount of EB1 comets/μm^2^ for non-rigor expressing cells (n = 105). Scalebars, 20μm.

To assess the effect of rigor-2mNeongreen expression on the subset of dynamic MTs, we expressed rigor-2xmNeongreen in U2OS cells and performed immunolabeling for end binding protein 1 (EB1). EB proteins bind to the tips of polymerizing MTs and are often used to visualize the subset of dynamic MTs (Akhmanova & Steinmetz 2015). To assess the effect of rigor-2xmNeongreen expression on dynamic MTs, we randomly imaged rigor- and non-rigor expressing cells. Subsequently, we counted the amount of EB1 comets per μm^2^ and measured the rigor-2xmNeongreen intensities per cell. We observed an inverse correlation between rigor-2xmNeongreen intensity and the density of EB1 comets, with a decrease in the density of EB1 comets with increasing rigor intensity (Fig.2C-D). Nonetheless, in low rigor expressing cells the amount of EB1 comets per μm^2^ was similar to control cells, indicating that the amount of polymerizing MTs in the cell is not affected by the presence of low levels of rigor-2xmNeongreen (Fig.2C-D). In conclusion, our data shows that, at low expression levels, rigor-2xmNeongreen induces minimal artefacts on the MT cytoskeleton.

Rigor-2xmNeongreen occupies the same binding sites on the MT as active kinesin-1. Therefore, expression of rigor-2xmNeongreen could potentially perturb endogenous kinesin-1-driven organelle transport. Distribution of mitochondria through the cell is kinesin-1 dependent and absence of kinesin-1 or its activating MAP, MAP7, leads to clustering of mitochondria around the nucleus (Hooikaas et al. 2019; Serra-Marques et al. 2020). To assess the effect of rigor-2xmNeongreen expression on kinesin-1-driven organelle transport, we scored mitochondrial spreading in random rigor- and non-expressing cells. In high expressing cells, mitochondria were tightly clustered around the nucleus in the majority of cells (67%) (Fig.3A,B). In addition, mitochondria often lost their long tubular structures and became fragmented (Sup.Fig.4A). However, in low expressing cells, mitochondria were spread through the cytoplasm in most cells (66.0%) which is similar to control cells (69.9%) (Fig.3A,B). Similar results were found in a duplicate experiment (Sup.Fig.4B). Next, we set out to show directly that organelle transport can take place along rigor-decorated MTs. Rab6a secretory vesicles are transported from the Golgi to the plasma membrane by concerted action of Kif5b and Kif13b (Serra-Marques et al. 2020). By simultaneous live-cell imaging of rigor-2xmNeongreen and rab6a using TIRF-microscopy, we could directly observe transport of rab6a vesicles along both straight and curvy parts of rigor-decorated MTs (3C-H). Taken together, these experiments indicate that at low expression levels, organelle transport can take place along rigor-decorated MTs.

**Figure 3.**
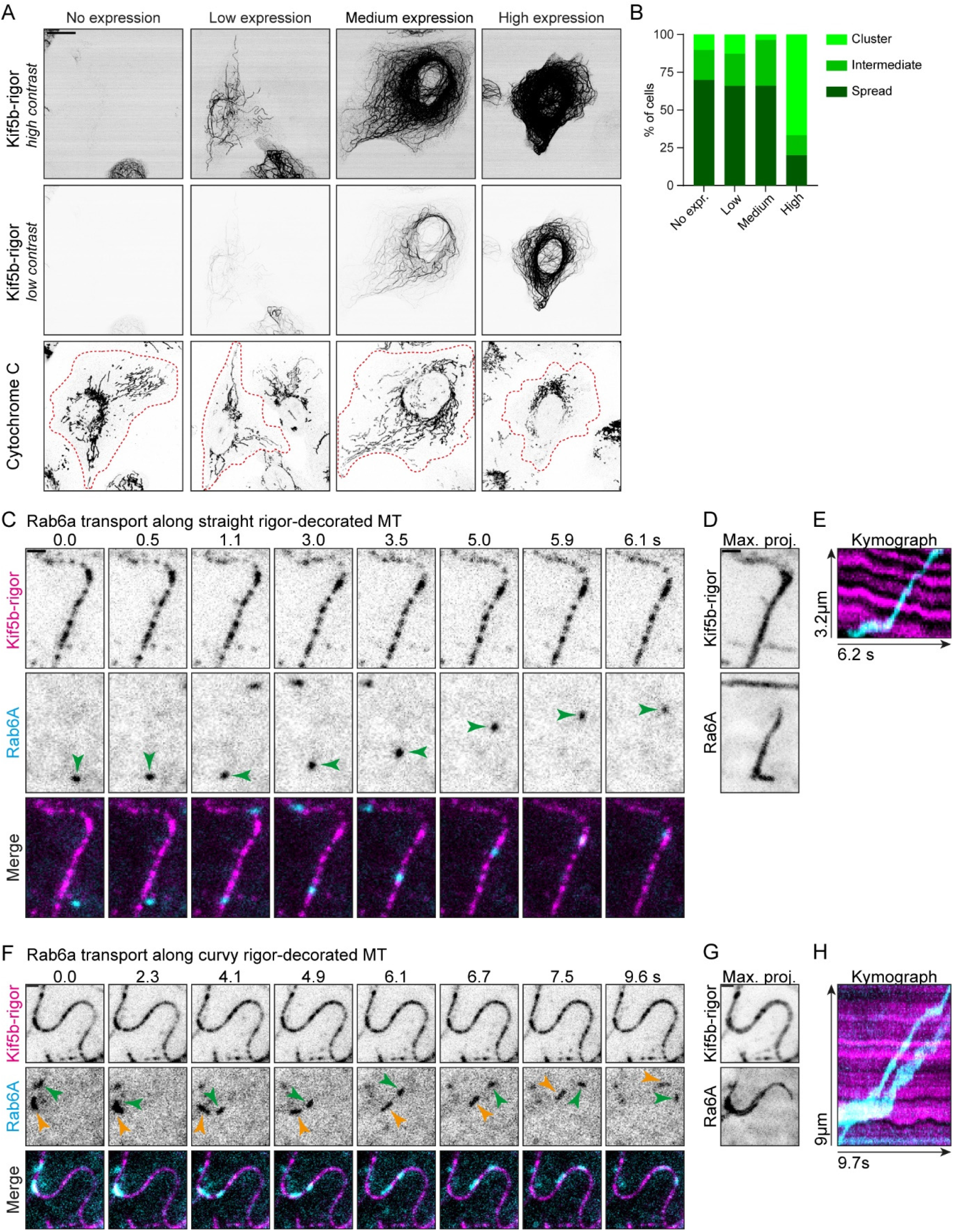
Rigor-decorated microtubules do not obstruct organelle transport. **(A)** Fluorescence images of U2OS cells stained for the mitochondrial marker cytochrome C at different levels of rigor-2xmNeongreen expression. Cell outlines are indicated with red dashed lines. **(B)** Classification of mitochondria distribution at different levels of rigor-2xmNeongreen expression (no expression, n = 166 cells; rigor-2xmNeongreen expressing, n = 119 cells (low, n = 47 cells; medium, n = 56 cells; high, n = 15 cells)) **(C,F)** Stills from live-cell imaging of U2OS cells expressing rigor-2xmNeongreen and mCherry-Rab6a (see also video 4 and 5). **(D,G)** Maximum projection over time from the stream acquisition shown in (C,F). **(E,H)** Kymograph showing Rab6a motility along rigor-decorated MT as in (C,F). Scalebars 20μm (A), 1μm (C,D,F,G). Time, seconds:milliseconds.

To examine if rigor-decorated MTs can still be disassembled, we performed a serum starvation assay. Earlier experiments have shown that the majority of stable MTs, identified by their nocodazole-resistance and immunolabelling for detyrosinated tubulin, disappear upon prolonged serum starvation and reappear again upon the addition of serum in 3T3 cells (Gundersen et al. 1994). To test whether rigor-decorated MTs respond to these physiological clues, we performed a serum starvation assay with Swiss 3T3 cells (Fig4A) and stained for acetylated tubulin as a marker for stable MTs, as this antibody was in our hands superior to the detyrosinated tubulin antibody. Control cells cultured in full medium have a clear population of stable MTs in the perinuclear region. Upon prolonged serum starvation, the majority of these MTs is lost and only a few remaining stable MTs can be observed. In cells subjected to prolonged serum starvation followed by 8hrs of full medium, a perinuclear network of stable MTs is re-established (Fig.4B). The same response to prolonged serum starvation and re-addition of serum could be observed in Swiss 3T3 cells expressing rigor-2xmNeongreen, where stable MTs are decorated by the rigor protein (Fig.4C). This indicates that the subset of stable MTs can still undergo changes and respond to physiological clues in presence of rigor-2xmNeongreen. These experiments demonstrate that acetylated, rigor-decorated MTs are not over-stabilized, as they largely disappear during prolonged serum starvation.

**Figure 4.**
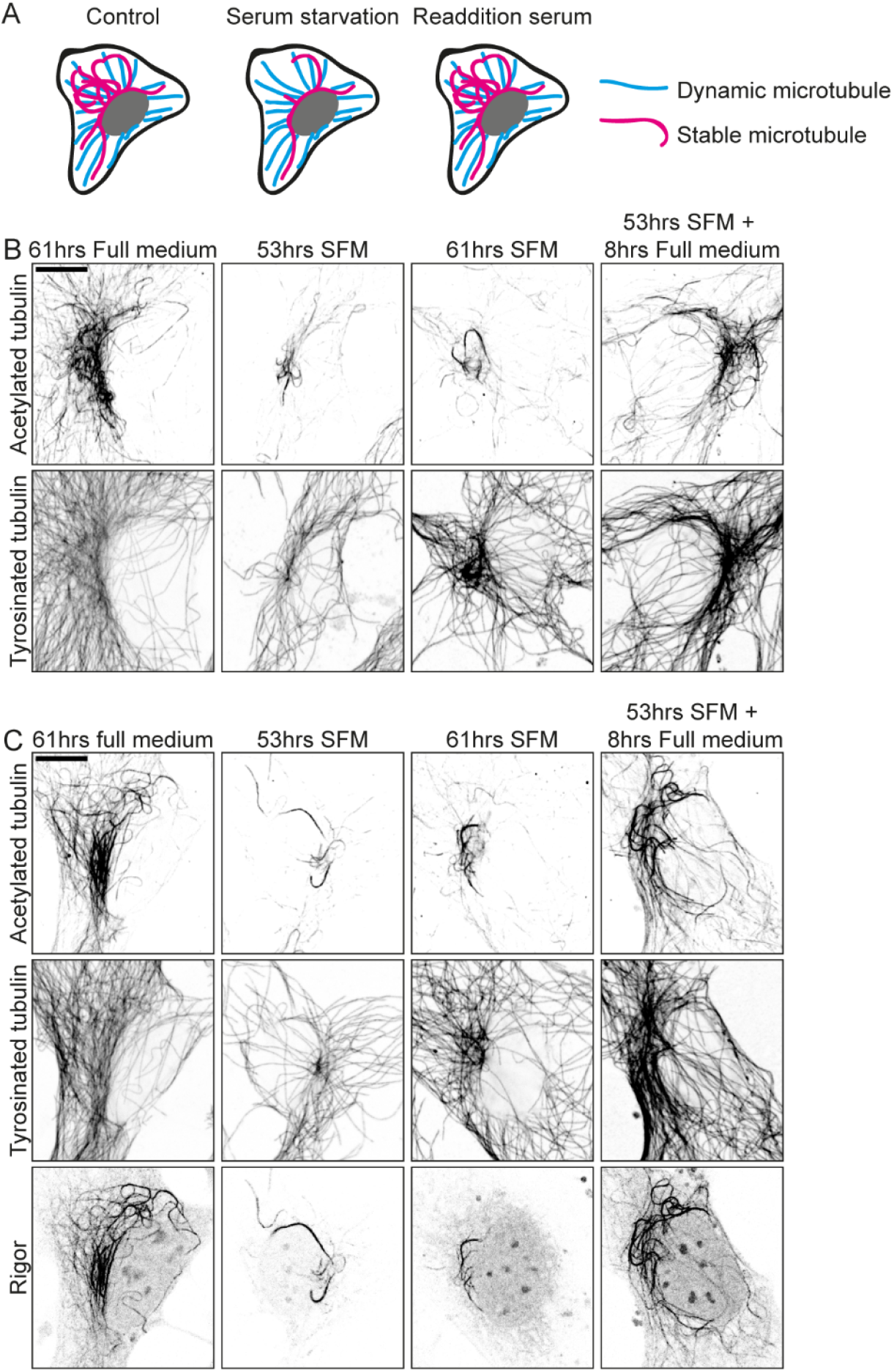
Rigor-decorated microtubules disappear upon serum starvation. **(A)** Cartoon serum starvation assay: under control conditions, Swiss 3T3 fibroblasts have a clear network of stable MTs. Upon prolonged serum starvation, the majority of these MTs is lost. A network of stable MTs re-emerges upon addition of serum. **(B)** Immunolabeling of acetylated- and tyrosinated MTs in Swiss 3T3 fibroblasts after 61hrs in full medium, 53hrs in serum-free medium (SFM), 61hrs in SFM or after 53hrs SFM followed by 8hrs in full medium. **(C)** Immunolabeling of acetylated- and tyrosinated MTs in Swiss 3T3 fibroblasts expressing rigor-2xmNeongreen under the same conditions as in (B). Experiment was performed in triplo – representative images are shown. Scalebars, 10μm.

Having validated rigor-2xmNeongreen as a live-cell marker for stable MTs, we set out to explore the dynamics of stable MTs in living cells using TIRF-microscopy. This revealed that, although these MTs are long-lived, they are far from static. Instead, stable MTs are very dynamic and display a variety of distinct behaviours. Under standard cell culture conditions, we frequently observed sliding events, where rigor-decorated MTs are transported through the cytoplasm with a speed of ~1μm/s, presumably by MT-bound motor proteins (Fig.5A,B). Furthermore, we often observed MT deformations (Fig.5C,D), where rigor-decorated MTs deform by what appears to be collisions with other cellular structures. This was different from undulation of rigor-decorated MTs (Fig.5E,F), where t MTs underwent fluctuations that seemed to arise from local, active pulling and pushing forces exerted on these MTs (Katrukha et al. 2017; Bicek et al. 2009; Kent et al. 2016; Pallavicini et al. 2014). Occasionally, we found rigor-decorated MTs in a loop configuration (Fig.5G-H), which implies that these MTs are exposed to considerable amounts of force. Infrequently, we observed a small piece of MT breaking off a rigor-decorated MT (Sup. Fig. 5A), indicating again that stable MTs are under mechanical stress. Sometimes we observe transport of small rigor fragments, remnants of depolymerization or breakage events, through the cell (Fig.5I,J). Although depolymerization events occur only sporadically, they can be visualized with our new live-cell marker (Sup. Fig.5B,C). Finally, rigor-MTs that were mostly static during the duration of imaging could also be found (Sup.Fig.5D).

**Figure 5.**
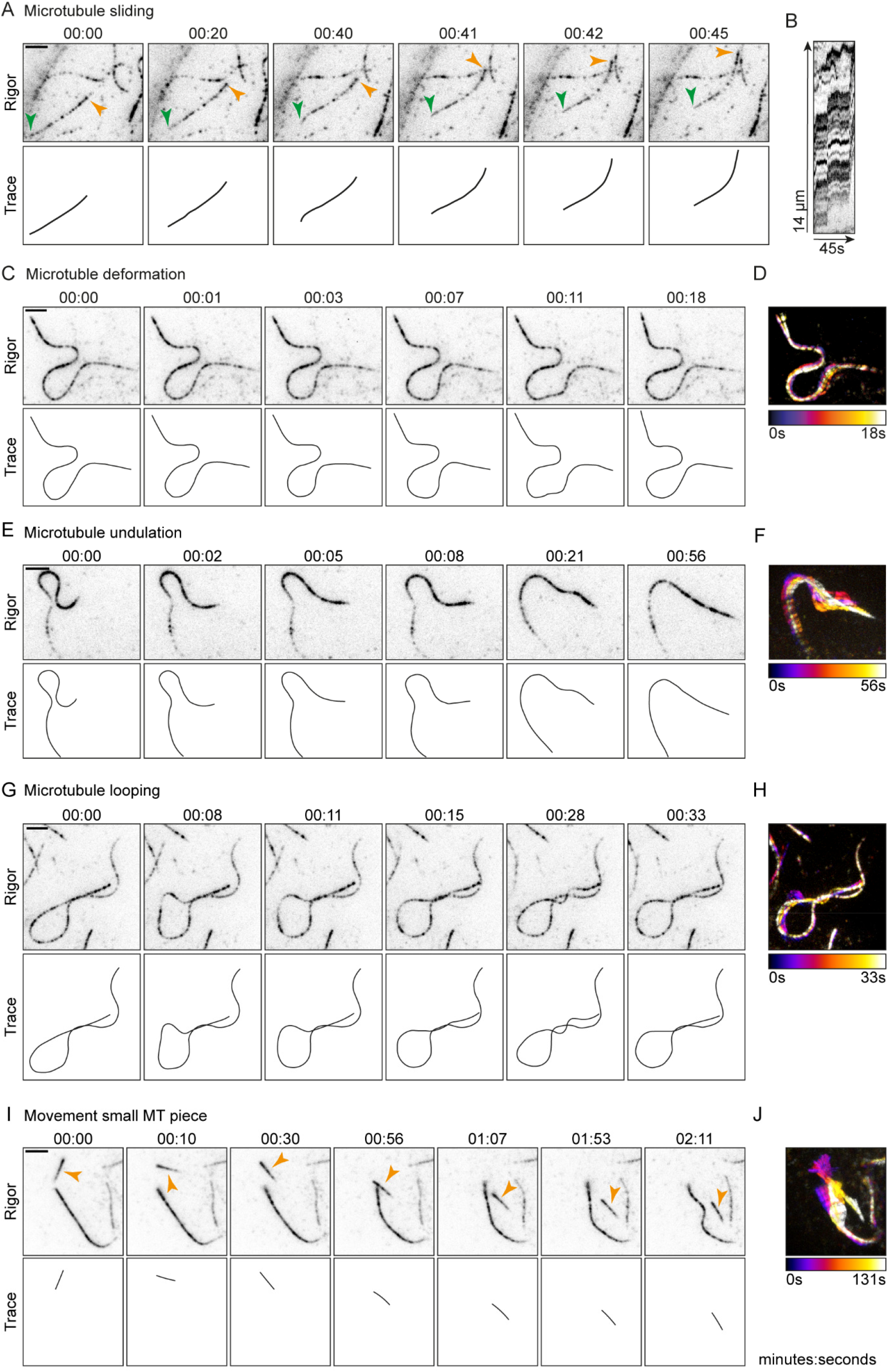
Stable microtubules undergo various forms of deformation. **(A,C,E,G, I)** Stills and schematic representations from live-cell imaging of rigor-2xmNeongreen in U2OS cells depicting MT sliding (indicated by coloured arrows), deformation, undulation, looping, and movement of a small MT piece (indicated by orange arrow), respectively (see also Video 6-10). **(B)** Kymograph of sliding event in (A). **(D, F, H, J)** Maximum projection of movie related to the still displayed in (C,E,G, I) color-coded for time. Scalebars, 2.5μm. Time, minutes:seconds.

While exploring the dynamics of stable MTs in living cells using TIRF-microscopy, we noticed very transient binding events of rigor molecules in regions of the cell devoid of long-lived MTs. We hypothesized that these could be binding events to dynamic MTs. To test if this was the case, we imaged U2OS cells expressing mCherry-tubulin and rigor-2xmNeongreen using fast TIRF-microscopy (5 frames/s) and found that rigor molecules can indeed bind transiently to dynamic MTs with an average dwell time of 0.65 s (Fig.6A-E). A maximum intensity projection of frames acquired in longer time lapses revealed MT shadows in the rigor channel in regions devoid of rigor-decorated, stable MTs (Fig.6F). These data indicate that rigor molecules randomly sample all MTs in the cell, but only become locked in their high affinity state when encountering a stable MT (Fig.6G), suggesting that the surface of stable MTs might be different from the surface of dynamic MTs.

**Figure 6.**
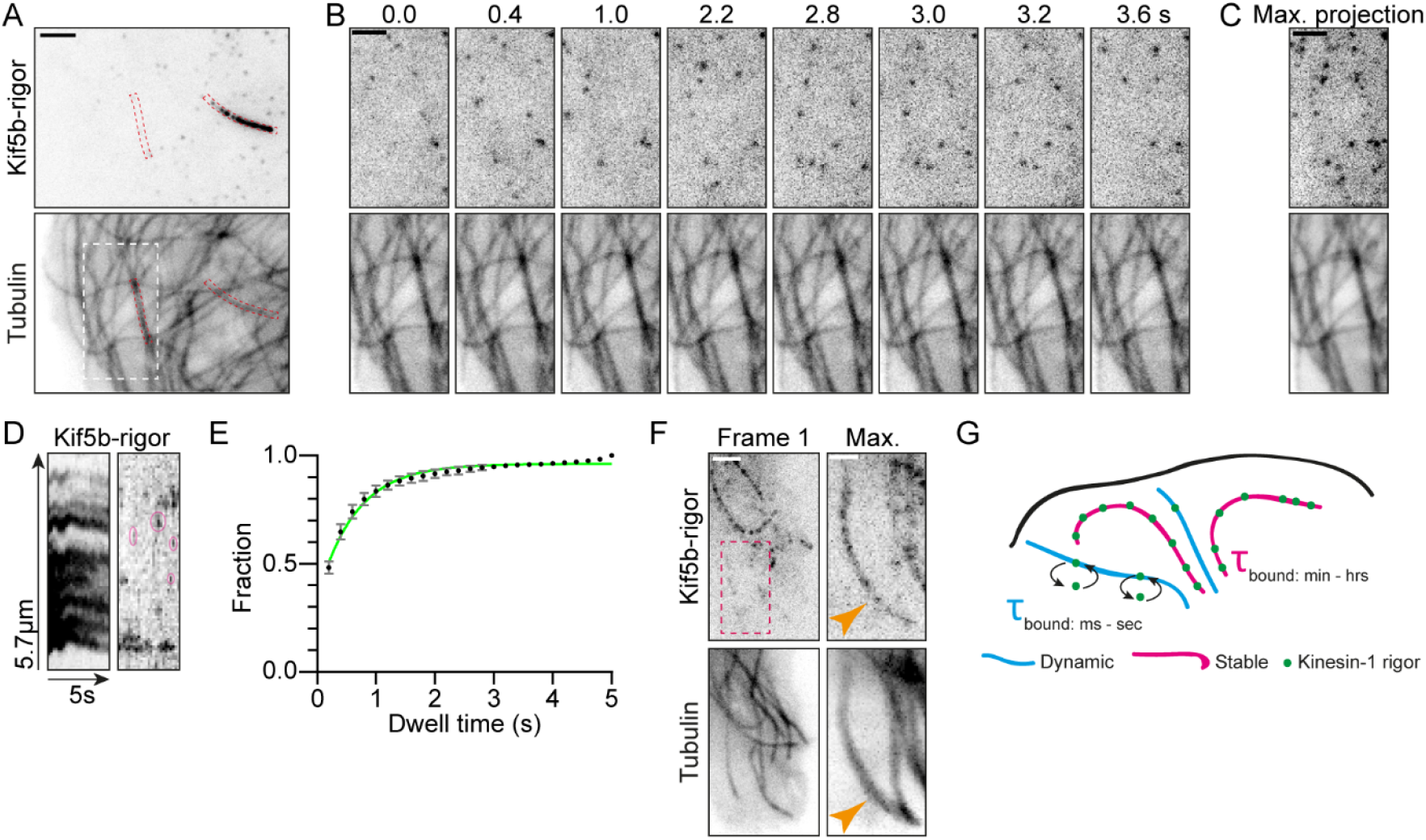
Rigor molecules bind transiently to dynamic microtubules. **(A)** Live U2OS cell expressing rigor-2xmNeongreen and mCherry-tubulin. **(B)** Stills from live-cell imaging of region indicated with white dashed box in (A) (see also video 11). **(C)** Maximum projection of 25 frames (5s) of region indicated with white dashed box in (A). **(D)** Kymographs of rigor channel of red dashed boxes in (A). Pink ovals in right kymograph indicate examples of short binding events of rigor to a dynamic MT. **(E)** Cumulative histogram + SD of dwell time of rigor on dynamic MTs (5006 events from 23 cells from 3 independent experiments). Green line shows fit with *y* = (*y*_1_ – *y*_2_)\**e*^(-*k*\**x*)^ + *y*_2_, yielding τ = 1/*k* = 0.65s **(F)** Live U2OS cell expressing rigor-2xmNeon and mCherry-tubulin. Maximum projection of red dashed box represents 123 frames (122s). Orange arrows point towards a dynamic MT with which rigor molecules have transiently interacted. **(G)** Cartoon: rigor molecules stably bind stable MTs (τ_bound_: minutes – hours), whereas rigor molecules interact transiently with dynamic MTs (τ_bound_: milliseconds – seconds). Scalebars 2.5μm (A), 2μm (B,C,E), 1μm (E – zoom). Time, seconds:milliseconds.

Finally, we further explored the mechanism by which stability is conferred to long-lived MTs. We considered two possible scenarios: (1) long-lived MTs are stabilized by mechanisms that prevent depolymerization of the plus-end; (2) long-lived MTs are stabilized along their whole length, e.g. by stabilizing MAPs or other modifications. To discriminate between these possibilities, we exploited laser-based severing of MTs. Previous laser-ablation experiments have revealed that upon severing of dynamic MTs, the freshly generated plus-end rapidly depolymerizes, whereas the minus-end often remains stable (Jiang et al. 2014). We reasoned that if the stability of long-lived MTs originates from their plus-end, the MT-ends generated by laser-based severing would still be susceptible to depolymerization. However, if long-lived MTs are stabilized along their whole length, freshly generated plus-ends would remain stable. To test this, we photo-ablated MTs in the perinuclear area of U2OS cells expressing mCherry-tubulin and rigor-2xmNeongreen using a high-power, pulsed laser (Fig.7A). Due to the high density of the MT network it was difficult to distinguish individual MTs labelled by mCherry-tubulin, but upon laser-induced severing we observed a wave of MT depolymerization occurring in the mCherry-tubulin channel, as expected for dynamic MTs. In contrast, MTs marked by rigor-2xmNeongreen largely remained stable on both sides of the cut (two examples shown in Fig.7B-C), i.e. 79% of the freshly generated ends showed no or little depolymerization (261 MT-ends from 36 cells from 4 independent experiments). We noted that also some rigor-negative MT-ends remained stable upon severing and we suspect that these correspond to minus-ends of dynamic MTs. Alternatively, these could indicate the existence of another, rigor-negative, subset of stable MTs. These results demonstrate that long-lived MTs are not just stabilized at their ends, but along their whole length.

**Figure 7.**
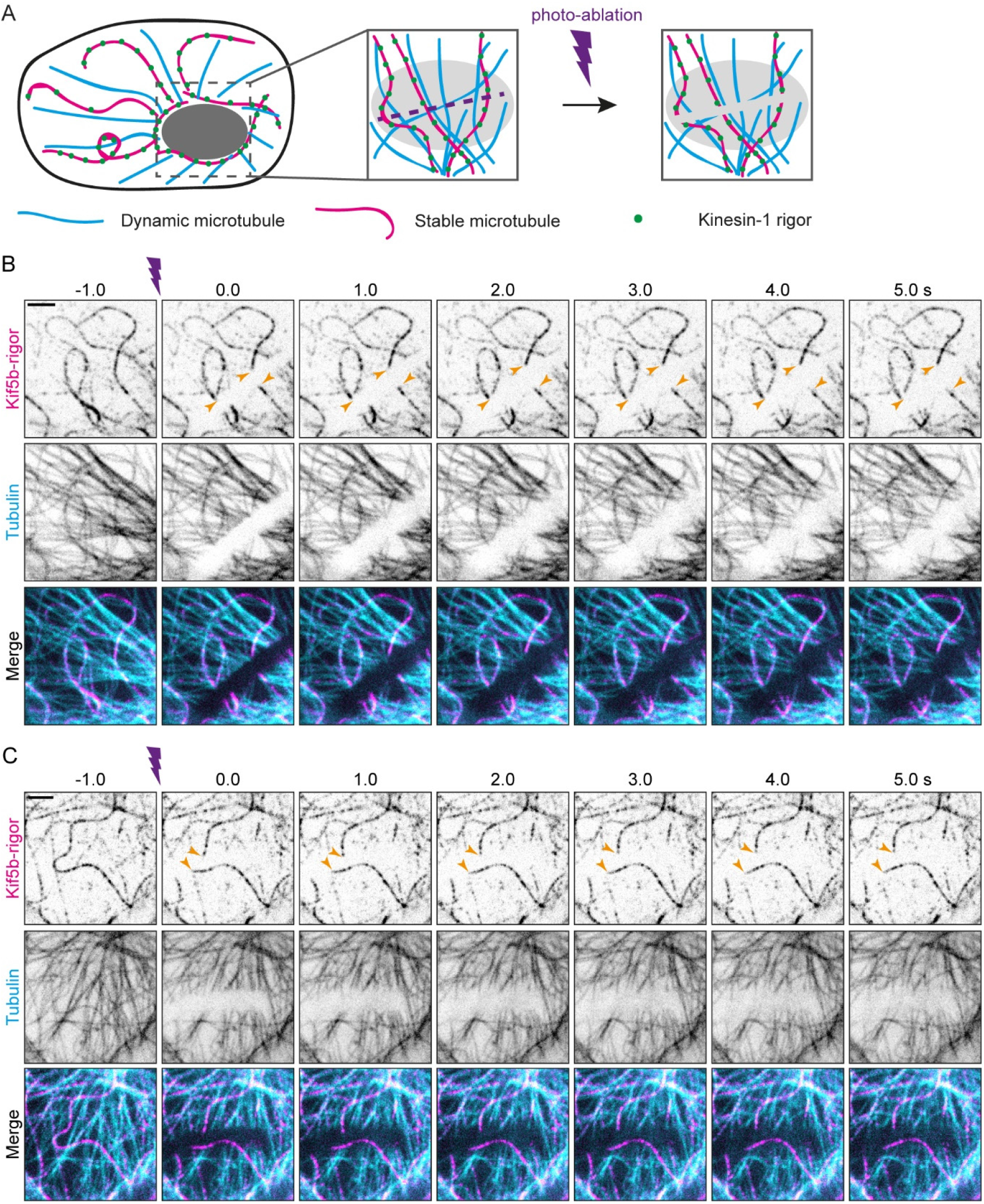
Laser-based severing of stable microtubules generates fresh MT-ends that are stable. **(A)** Cartoon: Stable and dynamic MTs in the nuclear area are severed by a focused laser beam. **(B-C)** Stills from live-cell imaging of U2OS cells expressing rigor-2xmNeongreen and mCherry-tubulin. Purple dart indicates photo-ablation. Coloured arrows indicate examples of rigor-decorated MT-ends that remain stable upon cutting (see also video 12-13). Scalebars, 2.5μm. Time, seconds:milliseconds.

## Discussion

Here, we present kinesin-1 rigor as a live-cell marker for stable MTs. Fusion to a tandem of the bright fluorophore mNeongreen (Shaner et al. 2013), allowed us to visualize low levels of kinesin-1 rigor bound to stable MTs in living cells. At these levels, expression of kinesin-1 rigor introduces minimal artefacts to the MT cytoskeleton, and organelle transport can still take place along rigor-decorated MTs. In addition, rigor-decorated MTs are responsive to external cellular clues; they disappear upon prolonged serum starvation and reappear upon addition of serum. Using our live-cell marker, we followed stable MTs in time and showed that these MTs are indeed long-lived. Furthermore, we show that on short timescales, stable MTs display very distinct behaviours. Finally, we provide evidence that long-lived MTs are stabilized by a mechanism that affects these MTs along their whole length.

Existing approaches to visualize stable MTs in living cells relied on imaging of a marker for total tubulin after nocodazole-treatment (Xu et al. 2017), which leaves the cells in very unphysiological conditions. Alternatively, researchers have used correlative live-cell and immunofluorescence microscopy (Cai et al. 2009), which is labour-intensive and error-prone given the continuous MT reorganization that we observed. The use of our live-cell marker is straightforward and allows direct visualization of the dynamics of stable MTs in the unperturbed cellular environment.

Our approach to visualize stable MTs in living cells relies on low levels of overexpression of a rigor mutant of kinesin-1. We characterized the effect of rigor expression on the MT cytoskeleton and find that the MT cytoskeleton is not altered by low levels of rigor expression. Yet we cannot completely exclude that low levels of our live-cell marker introduces minor artefacts on cellular physiology, as is the case with any cellular manipulation. Importantly, we find that, although kinesin-1 rigor and active kinesin-1 compete for the same binding sites on the MT, kinesin-1-driven organelle transport could still take place along rigor-decorated MTs. Indeed, *in vitro* work has shown that single kinesin-1 motors can make a transverse shift and circumvent kinesin-1-rigor road-blocks on the MT upon encounter (Schneider et al. 2015). Intracellular cargo is transported by a team of associated motor proteins. In line with our observations, it has been shown *in vitro* that a team of kinesins can overcome roadblocks and ensure smooth cargo transport (Tjioe et al. 2019; Ferro et al. 2019).

Fast imaging of stable MTs revealed a variety of dynamic behaviours, including sliding, deformation, undulation and looping. Kinesin-1-dependent MT sliding has been shown to drive changes in cell shape (Jolly et al. 2010) and is important for neurite outgrowth and axon regeneration in *Drosophila* (Lu et al. 2015; Del Castillo et al. 2015; del Castillo et al. 2015; Norkett et al. 2020; Winding et al. 2016) as well as cytoplasmic streaming in the *Drosophila* oocyte (Lu et al. 2016). Here, we find that MT sliding also takes place in interphase human cultured cells, indicating that sliding might be a phenomenon more common than originally thought and not exclusively a developmental process in specific cell types.

Undulation-like behaviours of MTs have been described before in *Xenopus laevis* melanophores (Pallavicini et al. 2014) and epithelial cells (Katrukha et al. 2017; Bicek et al. 2009; Kent et al. 2016). Recent work from our group demonstrated that these type of behaviours are not caused by contractions of the actomyosin network (Katrukha et al. 2017), but are likely caused by local force generators (Katrukha et al. 2017; Bicek et al. 2009; Kent et al. 2016; Pallavicini et al. 2014). A recent cryo-EM tomography study showed that MT curvature is higher in subcellular areas with high densities of actin- and intermediate filaments (Chakraborty et al. 2020). Nonetheless, the exact origin of these undulations-like motions is yet to be elucidated. Using our live-cell marker, we can now visualize these undulation-like behaviours of stable MTs with high spatiotemporal resolution, aiding future work aimed at understanding their origin.

TIRF-microscopy revealed that kinesin-1 rigor motors randomly bind to MTs of both subsets. However, rigor molecules quickly detach from dynamic MTs, while remaining stably bound to long-lived MTs. This suggests that the surface of stable MTs is different from the surface of dynamic MTs, and that this differentiation of the MT surface is ‘recognized’ by rigor motors, prompting them to adopt a high MT-affinity state. Our laser-ablation experiments also suggest that stable MTs have altered surface properties along their length compared to dynamic MTs. The exact mechanisms that could differentiate the surfaces of stable MTs from dynamic MTs are unknown, but could include PTMs, MT lattice decoration by different MAPs, incorporation of specific tubulin isotypes (Janke & Magiera 2020), or structural changes in the MT lattice itself.

Recent studies have started to unravel the importance of MT subsets in the spatial organization of especially complex cells like neurons (Burute & Kapitein 2019). However, these studies relied on detecting subsets in fixed cells and could not observe the differential dynamics of different MTs. We anticipate that the live-cell marker introduced in this work will help to understand how different MT subsets contribute to cellular organization and transport. In addition, it will help to unravel how complex MT arrays with multiple subsets are formed.

## Materials and Methods

### Cell culture and transfections

U2OS cells and Swiss 3T3 cells were cultured in DMEM supplement with 10% FCS and 50 μg/ml penicillin/streptomycin at 37°C and 5% CO_2_. U2OS cells were purchased from ATCC and Swiss 3T3 cells were a kind gift from Anna Akhmanova. Cells were confirmed to be free of mycoplasma. Cells were plated on 18mm coverslips (immunolabeling experiments) or 25mm coverslips (live-cell experiments) on the day of or 1 or 2 days before transfection. Cells were transfected using Fugene6 transfection reagent (Promega) according to manufacturer’s protocol. After 1 day of transfection, cells were used for live-cell imaging, subjected to treatment (serum starvation assay) or fixated. For all experiments that involved expression of mCherry-tubulin, cells were used for live-cell imaging after 2 days of transfection (Fig.1F-G, Fig.6A-E, Fig.7B-C, Sup.Fig.2A).

### Serum starvation assay

For the serum starvation assay, Swiss 3T3 fibroblasts were seeded on 18mm coverslips. The next day, cells were transfected with rigor-2xmNeongreen. 24hrs later, cells were washed 1X with starvation medium (DMEM supplemented with 50μg/ml penicillin/streptomycin) or full medium (DMEM supplement with 10% FCS and 50 μg/ml penicillin/streptomycin) and then incubated in starvation medium or full medium, respectively. 53hrs later, cells were washed 1X with starvation medium or full medium and were consecutively subjected to (a) fixation, (b) removal of starvation medium and addition of full medium, or (c) continuously incubated in fresh starvation medium or full medium. 8hrs later, cells were fixed.

### Plasmids and cloning

mCherry-α-tubulin (Kapitein et al. 2010) and FKBP-mCherry-Rab6A (Schlager et al. 2014) were described before. The presence of the N-terminal FKBP domain in the Rab6a construct has no detectable effects on the behaviour of this marker (Schlager et al. 2014; Serra-Marques et al. 2020).

hKif5b(1-560)-rigor-L-mNeongreen-L-mNeongreen was cloned into the mammalian expression vector pβactin-16-pl (chicken β-actin promoter) (Kaech et al. 1996) and generated by a combination of PCR-based cloning and Gibson assembly. The G234A rigor mutation was described previously (Rice et al. 1999). mNeongreen (Shaner et al. 2013) and flanking linker sequences were provided by Allele Biotechnology.

### Fluorescence microscopy

For live-cell imaging experiments, coverslips were mounted into metal imaging rings and immersed in full medium with (Fig.5A-J, Sup.Fig.5A-D) or without phenol red (Fig.1F-G, Fig.3C-D, Fig.6A-E, Fig.7A-B, Sup.Fig.2A). TIRF microscopy images (azimuthal spinning TIRF) were acquired on an inverted research Nikon Eclipse Ti-E microscope (Nikon) equipped with a perfect focus system (Nikon), ASI motorized stage MS-2000-XY (ASI), Apo TIRF 100x 1.49 NA oil objective (Nikon), iLas2 system (Roper Scientific, now Gataca system) and Evolve Delta 512 EMCCD camera (Photometrics). The microscope set-up was controlled by MetaMorph software 7.8 (Molecular Devices). For imaging, Stradus 488nm (150mW, Vortran) and OBIS 561nm (100mW, Coherent) lasers were used together with the ET-GFP filter set (49002, Chroma) or ET-GFP/mCherry filter set (59022, Chroma). Optosplit III beamsplitter (Cairn Research Ltd) equipped with a double emission filter cubed with ET525/50m, ET630/75m and T585LPXR (Chroma) was used during simultaneous imaging of green and red fluorescence. 16-bit images were projected onto the EMCCD chip with intermediate lens 2.5 X (Nikon C mount adapter 2.5X) at a magnification of 0.065 μm/pixel. To keep cells at 37°C, we used a stage top incubator (model INUBG2E-ZILCS, Tokai Hit). Images were acquired with 30 sec interval (Fig.1G), 1 sec interval (Fig.1F, Fig.5A-D, Fig.6E, Sup.Fig.5), at a frame rate of 5 (Fig.6A-D) or 10 frames/s (Fig.3C-D).

For the live nocodazole (10μM final concentration) (Cat#M1404, Sigma) assay, images were acquired with spinning disk confocal microscopy on inverted research microscope Nikon Eclipse Ti-E (Nikon), equipped with the perfect focus system (Nikon), Plan Fluor 40X 1.3 NA oil objective (Nikon) and a spinning disk-based confocal scanner unit (CSU-X1-A1, Yokogawa). The system was also equipped with ASI motorized stage with the piezo plate MS-2000-XYZ (ASI), Photometrics Evolve Delta 512 EMCCD camera (Photometrics) and controlled by MetaMorph 7.8 software (Molecular Devices). For imaging, Stradus 488nm (150mW, Vortran) and 561 Jive (100mW, Cobolt) lasers were used, together with ET-GFP (49002, Chroma) and ET-mCherry (49008, Chroma) filter sets. For simultaneous imaging of green and red fluorescence ET-GFP/mCherry (59022) filter set together with DV2 beamsplitter (Photometrics) were used. 16-bit images were projected onto the EMCCD chip with intermediate lens 2.0X (Edmund Optics) at a magnification of 0.164 μm/pixel. To keep cells at 37°C, we used stage top incubator (model INUBG2E-ZILCS, Tokai Hit). Images were acquired at 2.5 min. interval (Sup. Fig.2A). Video 1 was acquired on this set-up using a Plan Apo VC 60x N.A. 1.40 oil objective (Nikon). Images were acquired at 1.5 s interval at a magnification of 0.110 μm/pixel.

For the photo-ablation experiments, the same setup was used. Photo-ablation was performed with FRAP/PhotoAblation scanning system iLas (Roper Scientific France, now Gataca systems) mounted on custom-ordered illuminator (MEY10021; Nikon) and 355nm passively Q-switched pulsed laser (Teem Photonics) combined with S Fluor 100X N.A. 0.5-1.3 oil objective (Nikon). 16-bit images were projected onto the EMCCD chip at a magnification of 0.066 μm/pixel. Images were acquired at 2 frames/sec (Fig.7B-C).

Immunolabelled cell samples were acquired on a Zeiss Axio Observer Z1 LSM700 (Zeiss) using the Plan-apochromat 63X 1.4 NA oil DIC objective (Zeiss). For the acetylation, EB1 and cytochrome C control experiments, transfected and non-transfected cells from the same coverslip were randomly selected for imaging. Illumination settings were chosen so that low rigor expressing cells could be detected while at the same time no pixel saturation occurred for high expressing cells. Illumination settings were kept the same for all images acquired from a particular coverslip. For the fixed starvation assay, illumination settings were kept similar for all conditions.

### Immunofluorescence

For immunofluorescence experiments, different fixation methods were exploited depending on the antibodies used. For immunocytochemistry of MTs, cells were extracted for 1 min. in pre-warmed extraction buffer (0.3% Triton X-100 and 0.1% glutaraldehyde in BRB80 buffer (BRB80 buffer: 80mM Pipes, 1mM EGTA and 4mM MgCl2, pH 6.8) and subsequently fixed in pre-warmed 4% PFA in PBS for 10min. For immunocytochemistry of cytochrome C, cells were fixed in pre-warmed 4% PFA in PBS for 10min. For immunocytochemistry of EB1, cells were fixed in ice-cold methanol for 10min. After fixation, cells were washed with PBS, permeabilized with 0.25% Triton X-100 in PBS, washed again with PBS and subsequently blocked for 1hr with 3% BSA in PBS. Cells were incubated with primary antibody diluted in 3% BSA in PBS for 1hr at RT, washed with PBS and incubated with secondary antibody diluted in 3% BSA in PBS for 1hr at RT. After washing with PBS, cells were dipped in MQ, air-dried and mounted on microscopy slides using Prolong Diamond (Molecular Probes). The following primary antibodies were used in this study: Cytochrome C (6H2.B4, BD Biosciences), EB1 (5/EB1 BD Biosciences), acetylated tubulin (6-11B-1, Sigma-Aldrich), alpha-tubulin (EP1332Y, Abcam), alpha-tubulin (B-5-1-2, Sigma), tyrosinated tubulin (YL1/2, Abcam) and detyrosinated tubulin (AB3210, Merck).

### Image processing and analysis

To prepare images and movies for publication, FIJI was used to adjust contrast levels and perform background corrections, for bleach correction using histogram matching, to generate maximum intensity projections, and to generate kymographs using the FIJI-plugin KymoResliceWide (https://github.com/ekatrukha/KymoResliceWide). Chromatic correction of dual-colour live-cell imaging data was performed using the FIJI-plugin DoM_Utrecht (https://github.com/ekatrukha/DoM_Utrecht).

To assess the effect of rigor expression on microtubule acetylation levels, a ROI was drawn around every individual cell and the mean gray value (MGV) of every channel was measured. Data was transferred to Excel and the MGV of every channel was background corrected. For individual cells, the ratio between [MGV acetylated tubulin] / [MGV alpha-tubulin] was calculated. For rigor-expressing cells, the ratio between [MGV rigor] / [MGV alpha-tubulin] was also calculated.

To assess the effect of rigor expression on the amount EB1 comets, a ROI was drawn around individual cells and their surface area was measured. The amount of EB1 comets was counted using the ComDet plugin for FIJI (https://github.com/ekatrukha/ComDet). For rigor-expressing cells, the MGV of the rigor channel was measured. Data was transferred to Excel and for every cell the amount of EB1 comets per μm2 was calculated. MGVs of the rigor channel were background corrected.

To assess the effect of rigor expression on mitochondrial spreading, mitochondria distribution was classified as (a) spread throughout the cell, (b) clustered around the nucleus, or (c) intermediate phenotype where clustering around the nucleus was observed as well some spreading through the cytoplasm. During the classification of mitochondria distribution, the observer was unaware of the transfection status of the classified cell. For rigor-expressing cells, a ROI was drawn around the cell and MGV of the rigor channel was measured. Data was transferred to excel and MGVs of the rigor channel were background corrected. The resulting values were converted to Log10 values. Cells were classified based on their Log10(MGV) value into low, medium or high expression, with every class containing a range of values representing 1/3 of the difference between the Min and Max Log10 value of the dataset.

To assess the dwell time of rigor molecules bound to dynamic MTs, ROIs that were devoid of stable MTs and where binding-unbinding events were clearly visible were selected. Rigor molecules were automatically detected and fitted using the FIJI-plugin DoM_Utrecht (https://github.com/ekatrukha/DoM_Utrecht). Subsequently, to estimate the time rigor molecules stayed bound to dynamic MTs, detected molecules were linked in consecutive frames with a maximum distance of 4 pixels between detected particles in consecutive frames and a maximum linking gap in frames of 1 using the FIJI-plugin DoM_Utrecht. Data was transferred to excel and tracks lengths were multiplied by 0.2s to calculate binding times. Data was transferred to GraphPad Prism 9 to calculate cumulative frequency distributions. Subsequently, data was fitted with *y* = (*y*_1_ – *y*_2_)\**e*^(−*k*\**x*)^ + *y*_2_ to get the dwell time τ = 1/*k*. Data was plotted using GraphPad Prism 9.

## Acknowledgements

We thank Casper Hoogenraad for the mCherry-α-tubulin and FKBP-mCherry-Rab6A constructs and Anna Akhmanova for fruitful discussion and for providing the Swiss 3T3 cells. This work is supported by the Netherlands Organization for Scientific Research (NWO; NWO-Graduate program project 022.006.001 to K.I. Jansen), European Molecular Biology Organization (EMBO; EMBO long-term fellowship (EMBO ALTF 407-2017) to M. Burute) and the European Research Council (ERC Consolidator Grant 819219 to L.C. Kapitein).

The authors declare no competing financial interests.

## Author contributions

K.I. Jansen created reagents, performed experiments, and analysed data. M.B. performed additional experiments. K.I. Jansen and L.C. Kapitein designed the study, interpreted data, and wrote the manuscript. L.C. Kapitein supervised the project.

---

**Supplemental Figure 1.**
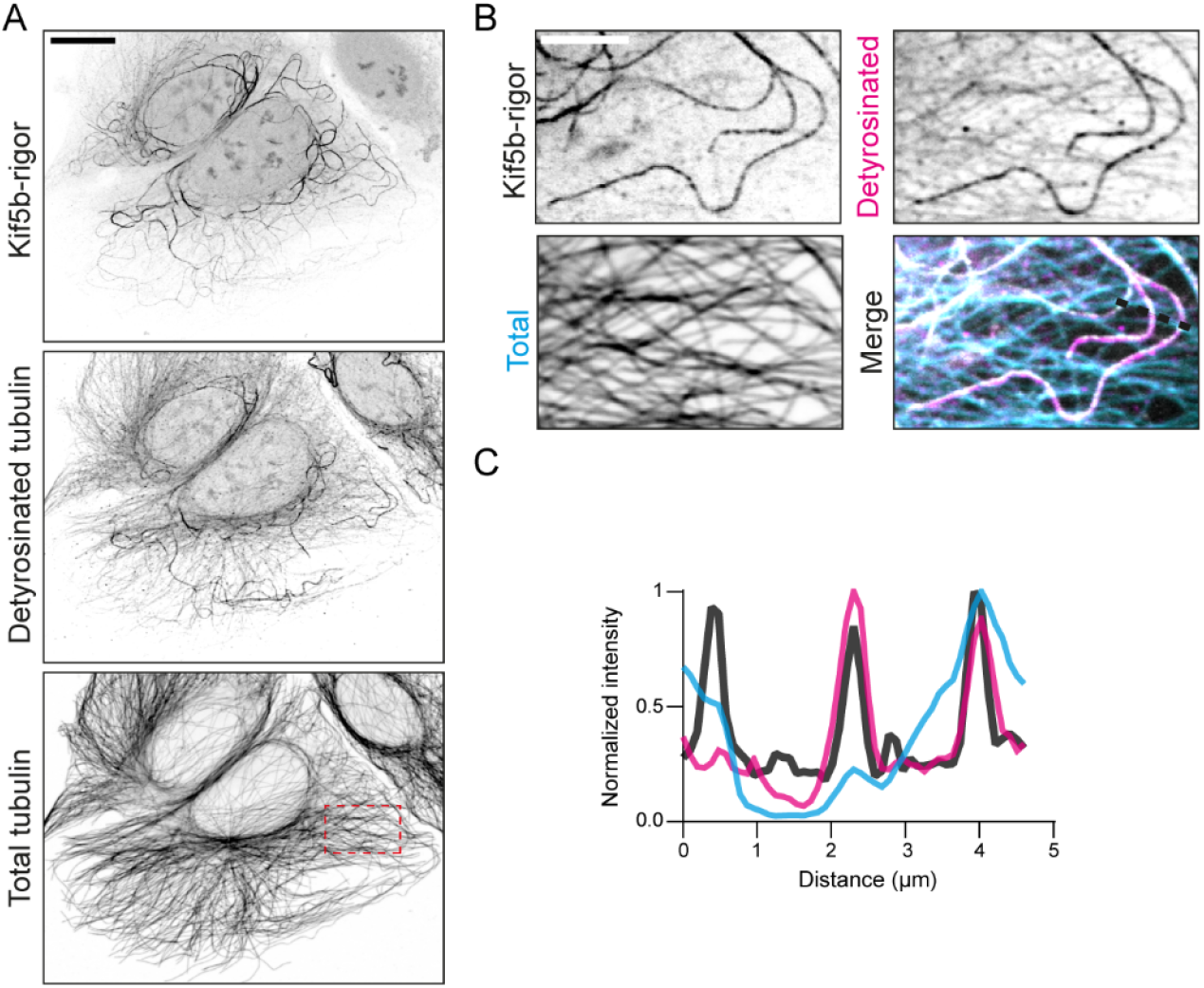
related to Fig.1. Kinesin-1 rigor decorates detyrosinated microtubules. **(A)** Fluorescence images of U2OS cell expressing rigor-2xmNeongreen and immunolabelled for detyrosinated tubulin and alpha-tubulin shown in inverted contrast. **(B)** Zooms of region indicated by dashed box in (A). **(C)** Intensity profile across the region indicated with dashed line in (B). Scalebars, 15μm (A), 5μm (B).

**Supplemental Figure 2.**
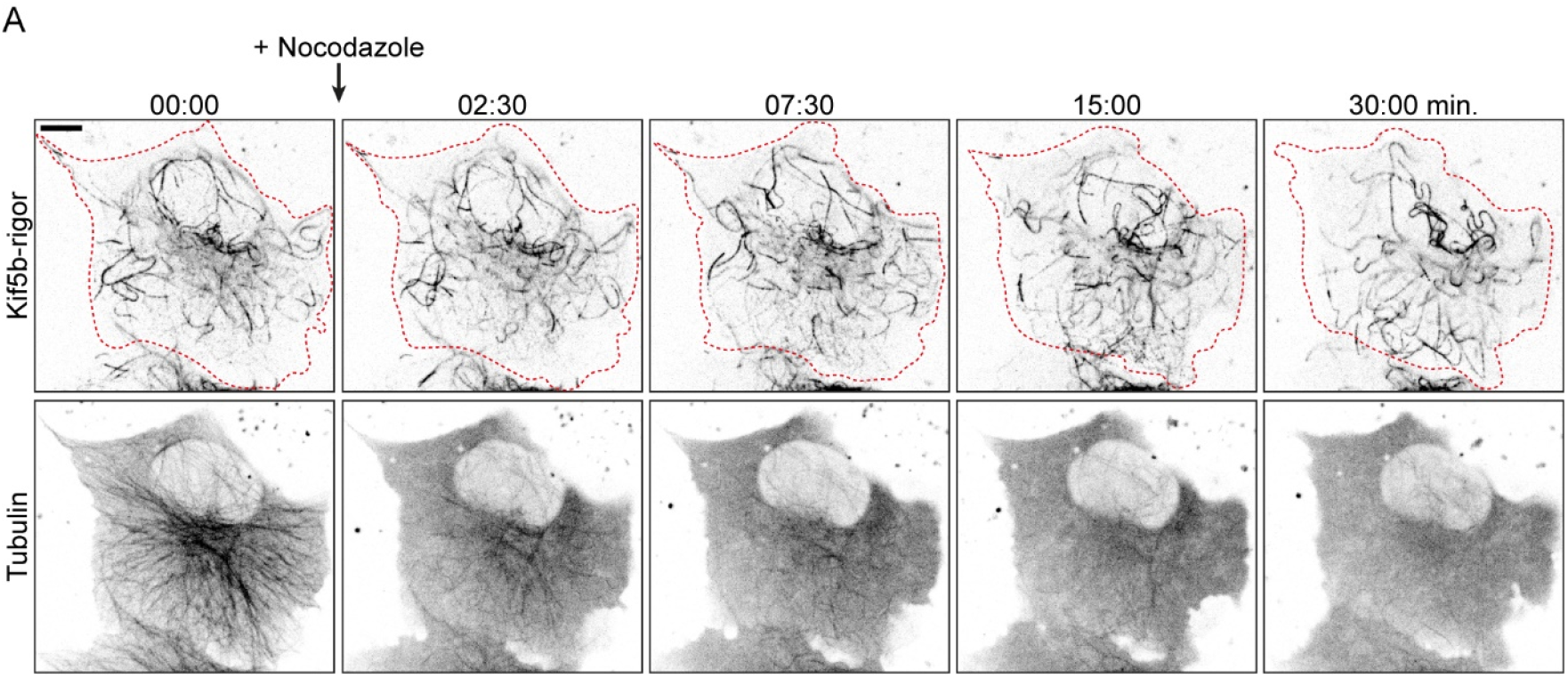
related to Fig.1. Rigor-decorated microtubules are resistant to nocodazole-induced depolymerization. (A) Stills from live-cell imaging of rigor-2xmNeongreen and mCherry-tubulin in U2OS cell upon addition of 10μM nocodazole (see also Video 14). Scalebar, 10μm. Time: minutes:seconds.

**Supplemental Figure 3.**
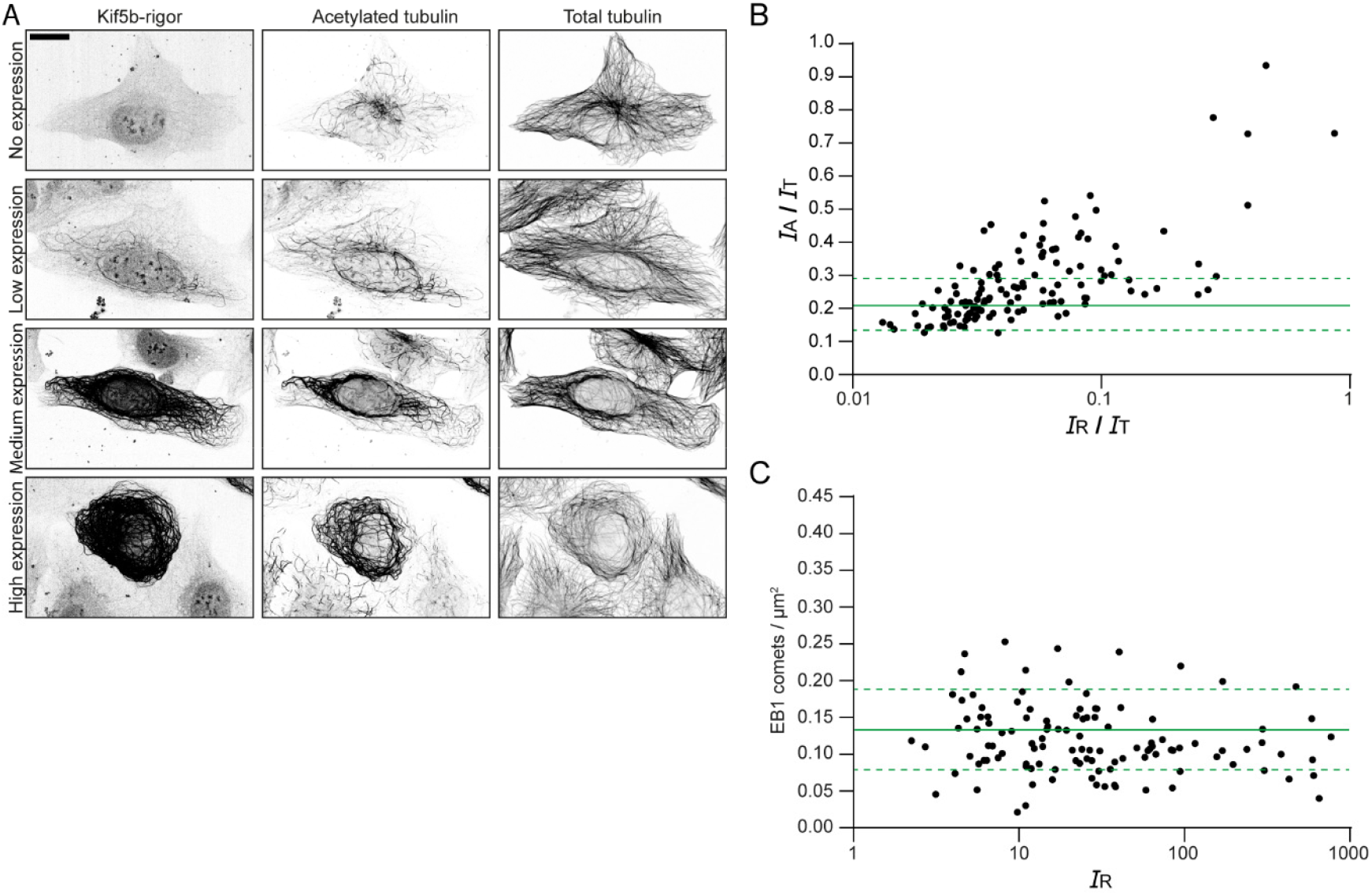
related to Fig. 2 Kinesin-1 rigor has minimal effects on the microtubules cytoskeleton at low levels. **(A)**Fluorescence images of U2OS cells stained for acetylated tubulin and alpha-tubulin at different levels of rigor-2xmNeongreen expression as analysed in (B). **(B)** Quantification showing the intensity ratio of acetylated tubulin (*I*_A_) over total tubulin (*I*_T_), plotted against the intensity ratio of rigor-2xmNeon (*I*_R_) over total tubulin for individual rigor-expressing cells (each dot represents a single cell, n = 131). Solid green line + dashed lines indicate mean ± SD of the intensity ratio of acetylated tubulin over total tubulin for non-rigor expressing cells (n = 120). **(C)** Quantification showing the amount of EB1 comets/μm^2^ for cells with different rigor-2xmNeongreen intensities (each dot represents a single cell, n = 115). Solid green line + dashed lines indicated mean + SD of amount of EB1 comets/μm^2^ for non-rigor expressing cells (n = 99). Scalebars, 20μm.

**Supplemental Figure 4.**
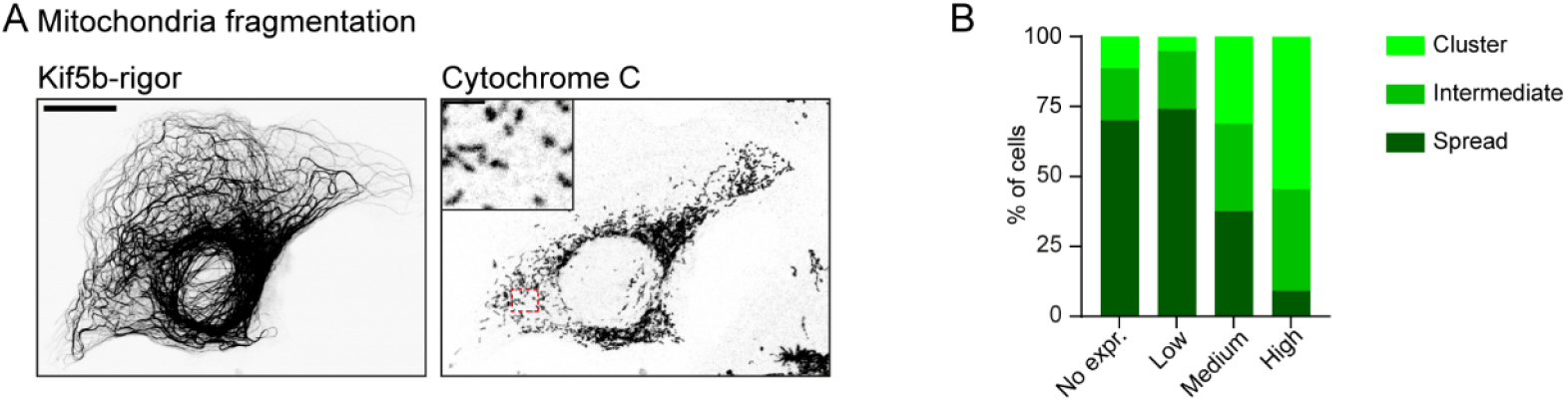
related to Fig. 3 Rigor-decorated microtubules do not obstruct organelle transport. **(A)** Fluorescence images of U2OS cell stained for the mitochondrial marker cytochrome C at high level of rigor-2xmNeongreen expression. Inset displays zoom of the region indicated with red box. **(B)** Classification of mitochondria distribution at different levels of rigor-2xmNeongreen expression (no expression, n = 124 cells; rigor-2xmNeongreen expressing, n = 101 cells (low, n = 58 cells; medium, n = 32 cells; high, n = 11 cells)) Scalebars, 20μm; zoom 2.5μm.

**Supplemental Figure 5.**
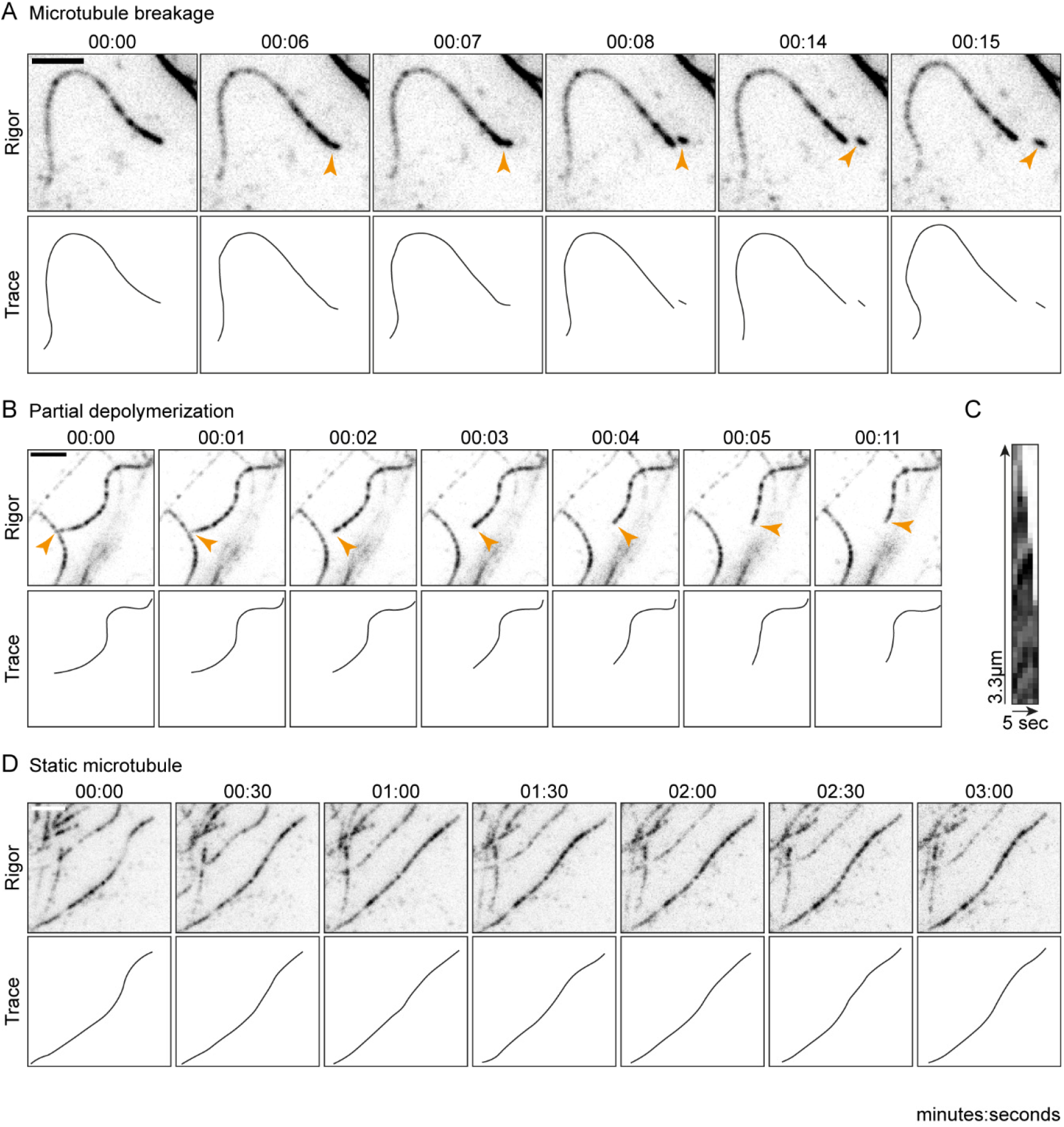
related to Fig. 5 Stable microtubules display different types of motility. **(A,B,D)** Stills and schematic representations from live-cell imaging of rigor-2xmNeongreen in U2OS cells depicting MT breakage (indicated by orange arrow), partial MT depolymerization (indicated by orange arrow) and a static MT, respectively (see also Video 15-17). **(C)** Kymograph of the partial depolymerization event shown in (B). Scalebars, 2.5μm. Time, minutes:seconds.

**Video 1 (related to Fig.1)**

Rigor-2xmNeongreen and mCherry-tubulin in U2OS cell. This video corresponds to Fig.1. Total time: 3min. Acquired with 1.5 s interval between frames. 13x sped up.

**Video 2 (related to Fig.1F)**

Rigor-2xmNeongreen and mCherry-tubulin in a U2OS cell. This video corresponds to Fig.1F. Total time: 1 min and 10 s. Acquired with 1 s interval between frames. 20x sped up.

**Video 3 (related to Fig.1G)**

Rigor-2xmNeongreen and mCherry-tubulin in a U2OS cell. This video corresponds to Fig.1G. Total time: 30min. Acquired with 30 s interval between frames. 225x sped up.

**Video 4 (related to Fig. 3C)**

Rigor-2xmNeongreen and mCherry-Rab6a in a U2OS cell. This video corresponds to Fig.3C. Total time: 6.2 s. Stream acquisition acquired at a speed of 10 frames/s. 2x sped up.

**Video 5 (related to Fig.3F)**

Rigor-2xmNeongreen and mCherry-Rab6a in a U2OS cell. This video corresponds to Fig.3F. Total time: 9.7 s. Stream acquisition acquired at a speed of 10 frames/s. 2x sped up.

**Video 6 (related to Fig.5A)**

Rigor-2xmNeongreen in U2OS cell. This video corresponds to Fig.5A. Total time: 45 s. Acquired with 1 s interval between frames. 10x sped up.

**Video 7 (related to Fig.5C)**

Rigor-2xmNeongreen in U2OS cell. This video corresponds to Fig.5C. Total time: 18 s. Acquired with 1 s interval between frames. 10x sped up.

**Video 8 (related to Fig.5E)**

Rigor-2xmNeongreen in U2OS cell. This video corresponds to Fig.5E. Total time: 56 s. Acquired with 1 s interval between frames. 10x sped up.

**Video 9 (related to Fig.5G)**

Rigor-2xmNeongreen in U2OS cell. This video corresponds to Fig.5G. Total time: 33 s. Acquired with 1 s interval between frames. 10x sped up.

**Video 10 (related to Fig.5I)**

Rigor-2xmNeongreen in U2OS cell. This video corresponds to Fig.5I. Total time: 2min and 11 s. Acquired with 1 s interval between frames. 10x sped up.

**Video 11 (related to Fig.6B)**

Rigor-2xmNeongreen and mCherry-tubulin in U2OS cell. This video corresponds to Fig.6B. Total time: 5 s. Stream acquired at 5 frames/s. 6x sped up.

**Video 12 (related to Fig.7B)**

Rigor-2xmNeongreen and mCherry-tubulin in U2OS cell. This video corresponds to Fig.7B. Total time: 6 s. Acquired at a frame rate of 2 frames/s. 5x sped up.

**Video 13 (related to Fig.7C)**

Rigor-2xmNeongreen and mCherry-tubulin in U2OS cell. This video corresponds to Fig.7C. Total time: 6 s. Acquired at a frame rate of 2 frames/s. 5x sped up.

**Video 14 (related to Sup. Fig.2A)**

Rigor-2xmNeongreen and mCherry-tubulin in U2OS cell. This video corresponds to Sup.Fig.2A. Total time: 30min. Acquired with 2.5min. interval between frames. 1125x sped up.

**Video 15 (related to Sup. Fig.5A)**

Rigor-2xmNeongreen in U2OS cell. This video corresponds to Sup. Fig.5A. Total time: 15 s. Acquired with 1 s interval between frames. 5x sped up.

**Video 16 (related to Sup. Fig.5B)**

Rigor-2xmNeongreen in U2OS cell. This video corresponds to Sup. Fig.5B. Total time: 11 s. Acquired with 1 s interval between frames. 10x sped up.

**Video 17 (related to Sup. Fig.5D)**

Rigor-2xmNeongreen in U2OS cell. This video corresponds to Sup. Fig.5D. Total time: 3 min. Acquired with 1 s interval between frames. 30x sped up.

## Notes

### Competing Interest Statement

The authors have declared no competing interest.

